# Dual-wavelength stopped-flow analysis of the lateral and longitudinal assembly kinetics of vimentin

**DOI:** 10.1101/2022.05.05.490764

**Authors:** Lovis Schween, Norbert Mücke, Stéphanie Portet, Wolfgang H. Goldmann, Harald Herrmann, Ben Fabry

## Abstract

Vimentin is a highly charged intermediate filament protein that inherently forms extended dimeric coiled-coils, which serve as the basic building blocks of intermediate filaments. Under low ionic strength conditions, vimentin filaments dissociate into uniform tetrameric complexes of two anti-parallel oriented, half-staggered coiled-coil dimers. By addition of salt, vimentin tetramers spontaneously reassemble into filaments in a time-dependent process: i) lateral assembly of tetramers into unit-length filaments (ULFs); ii) longitudinal annealing of ULFs; iii) longitudinal assembly of filaments coupled with subsequent radial compaction. To independently determine the lateral and longitudinal assembly kinetics, we measure with a stopped-flow instrument the static light scattering signal at two different wavelengths (405 and 594 nm) with a temporal resolution of 3 ms, and analyze the signals based on Rayleigh-Gans theory. This theory considers that the intensity of the scattered light depends not only on the molecular weight of the scattering object but also on its shape. This shape-dependence is more pronounced at shorter wavelengths, allowing us to decompose the scattered light signal into its components arising from lateral and longitudinal filament assembly. We demonstrate that both the lateral and longitudinal filament assembly kinetics increase with salt concentration.

**Significance statement:** The proper formation of intermediate filament (IF) networks in the cytoplasm is important for numerous cell functions. Here, we present a stopped-flow method for measuring the in-vitro assembly kinetics of intermediate filaments with a temporal resolution of 3 ms using static light scattering at two different wavelengths. This allows us to compute the shape factor of the assembly products based on Rayleigh-Gans light scattering theory. From the shape factor, we can separately measure the lateral assembly of tetramers into unit-length filaments (ULFs), and the longitudinal annealing of ULFs and longer filaments. For the IF protein vimentin, we find that with increasing salt concentrations, both the lateral and longitudinal assembly rates increase, and unstable, hyper-aggregated assembly complexes emerge.

## Introduction

Intermediate filament (IF) proteins constitute a central structural element in metazoan cells. IFs form, together with the F-actin and microtubule system, cell type-specific networks referred to as the cytoskeleton (1, 2). IFs withstand standard disruptive treatments during isolation from tissues and can be recovered from the insoluble fractions. IFs can then be solubilized by treatment with chaotropic reagents such as urea or by resuspension in buffer of very low ionic strength (3). Under low ionic strength conditions, IF proteins are present in form of tetrameric complexes, as revealed by analytical ultracentrifugation (4–7). By raising the ionic strength, tetramers spontaneously aggregate laterally in register and form so-called unit-length filaments (ULFs). Subsequently, ULFs anneal longitudinally into long filaments. The formation of ULFs from tetramers via octamers and 16-mers is fast, with rate constants >100 *μM*^−1^*s*^−1^. This process is largely completed within 1 s, as shown by light scattering measurements with a stopped-flow device (8). When the mass per cross-section is measured after 2 seconds of assembly using scanning transmission electron microscopy (STEM), the formed ULFs are distinctly heterogeneous, with values of 30 to 90 kDa/nm. This indicates that the tetramers, through various permutations of intermediate complexes such as octamers, 16-mers, and 24-mers, have assembled into higher-order complexes (8, 9). After 60 min of assembly, however, long and comparatively uniform filaments of 8 to 12 tetramers (38 to 56 kDa/nm) per cross-section prevail (9). This indicates that the complexes formed in the very first phase of assembly are in a dynamic transition state with respect to the lateral packing of tetramers, eventually reaching a stable number of tetramers per cross-section as weakly associated subcomplexes dissociate.

To gain deeper insights into the initial phase of filament elongation, we have implemented a stopped-flow device equipped with two lasers of different wavelengths, 405 nm and 594 nm, for static light scattering measurements with a time resolution of 3 ms. By measuring the scattered light at two different wavelength, we can disentangle the lateral and longitudinal assembly kinetics of vimentin based on a simple principle: during the early phase of the assembly process when the tetramers are laterally aggregating into ULFs, the static light scattering signal at both wavelengths increases in proportion with the molecular weight of the reaction products. When filaments elongate by longitudinal annealing of ULFs, however, the scattered light increases less-than-proportional with filament length, until no further significant increase in the scattered light intensity occurs. This is the case when filaments have grown beyond a length of ~500 nm (11 ULFs) for incident light with a wavelength of 594 nm, or ~250 nm (5-6 ULFs) for incident light with a wavelength of 405 nm.

The wavelength and filament length-dependence of the scattered light intensity can be described by the Rayleigh-Gans scattering theory. In particular, this theory allows us to compute the shape of the scattering objects and thus the longitudinal assembly rate from the ratio of the scattered light intensities at 594 nm and 405 nm. By contrast, lateral filament assembly equally affects the scattered light signals at different wavelengths and hence does not affect the intensity ratio. Thus, using a dual-wavelength stopped-flow approach, we can separately monitor the lateral and longitudinal components of the filament assembly process.

Our data reveal that with increasing salt concentrations, both the lateral and longitudinal assembly rates of vimentin increase, and the mature filaments exhibit a higher mass per cross-section. Intriguingly, we identify a salt and protein concentration-dependent transient reorganization phase (1-90 s after assembly start), during which higher-order complexes appear that are not found in mature filaments.

## Materials and Methods

### A. Protein chemical methods

Human vimentin is expressed in transformed bacteria, E. coli strain TG1, and isolated from inclusion bodies as described previously (3). For IF assembly, we use a Tris-HCl-buffered sodium chloride system. In brief, tetramers are re-natured by dialysis of monomers dissolved in 8M urea with 10 mM Tris-HCl (pH 7.5) against a solution containing 5mM Tris-HCl (pH 8.4) and 1 mM DTT. For the last dialysis step, the buffer is thoroughly degassed to prevent the formation of gas bubbles in the stopped-flow reaction chamber (3, 8, 10).

Assembly is initiated by addition of an equal volume of degassed salt buffer, i.e., low salt buffer: 45 mM Tris-HCl (pH 7.0) with 100 mM NaCl; medium salt buffer: 45 mM Tris-HCl (pH 7.0) with 200mM NaCl; high salt buffer: 45 mM Tris-HCl (pH 7.0) with 320 mM NaCl. Hence, the ionic conditions in the assembly chamber are 1.) low salt buffer 22.5 mM Tris-HCl (pH 7.5), 50 mM NaCl; 2.) medium salt buffer 22.5mM Tris-HCl (pH 7.5), 100mM NaCl; high salt buffer 22.5mM Tris-HCl (pH 7.5), 160mM NaCl. Final protein concentrations for assembly are between 0.05 – 0.4 mg/ml (corresponding to 0.25-2 *μM*).

### B. Atomic force microscopy

Atomic force microscopy (AFM) is performed using the tapping-mode in air as described previously (8, 10). Vimentin is assembled at 37°C in low salt buffer at a protein concentration of 0.4 mg/ml. Assembly is stopped by 10-fold dilution with low salt buffer followed by fixation with an equal volume of 0.2% glutaraldehyde in low salt buffer for 1 min. 40 *μl* are then deposited on freshly cleaved mica. After 1 min, the mica samples are washed with double-distilled water and dried with a steady stream of nitrogen.

### C. Transmission Electron Microscopy (TEM)

Samples for visualization by TEM are fixed with freshly prepared 0.2% glutaraldehyde in assembly buffer. The filament suspension is briefly adsorbed to glow-discharged carbon-coated electron microscopic grids, washed with distilled water, and negatively stained with 2% uranyl acetate for 15 seconds (3). Specimens are examined in a Zeiss (model 910, Carl Zeiss, Oberkochen, Germany) transmission electron microscope.

The filament width is determined near the center of the filaments using the image analysis software Clickpoints (11). The diameters of at least 77 filaments for each time point (2, 10, 30, 60, 180, 300 s after assembly start) are measured.

### D. Immunofluorescence Microscopy

The filament suspension is absorbed on ethanol-cleaned glass slides for 1 min followed by fixation in methanol (6 min) and acetone (30 s) at −20°C. Primary rabbit anti-vimentin antibodies (12) and fluorescently labeled secondary antibodies (Jackson laboratories) are used to visualize the fixed filaments (13).

### E. Stopped-flow experiments and data management

Stopped-flow measurements are performed with a two-syringe stopped-flow apparatus (SF-61, Hi-Tech Scientific, Salisbury, UK) as described in detail in (8) (Fig. 1). In brief, one syringe is filled with soluble vimentin protein (concentration between 0.1–0.8 mg/ml), and the other syringe with the assembly buffer containing 45 mM Tris-HCl, 100-320 mM NaCl at pH 7.0. All solutions are filtered through 0.22 *μl* syringe filters (Rotilabo; Carl Roth, Karlsruhe, Germany). Both syringes are heated to 37°C and allowed to equilibrate for 5 min before starting the measurements.

**Fig. 1.**
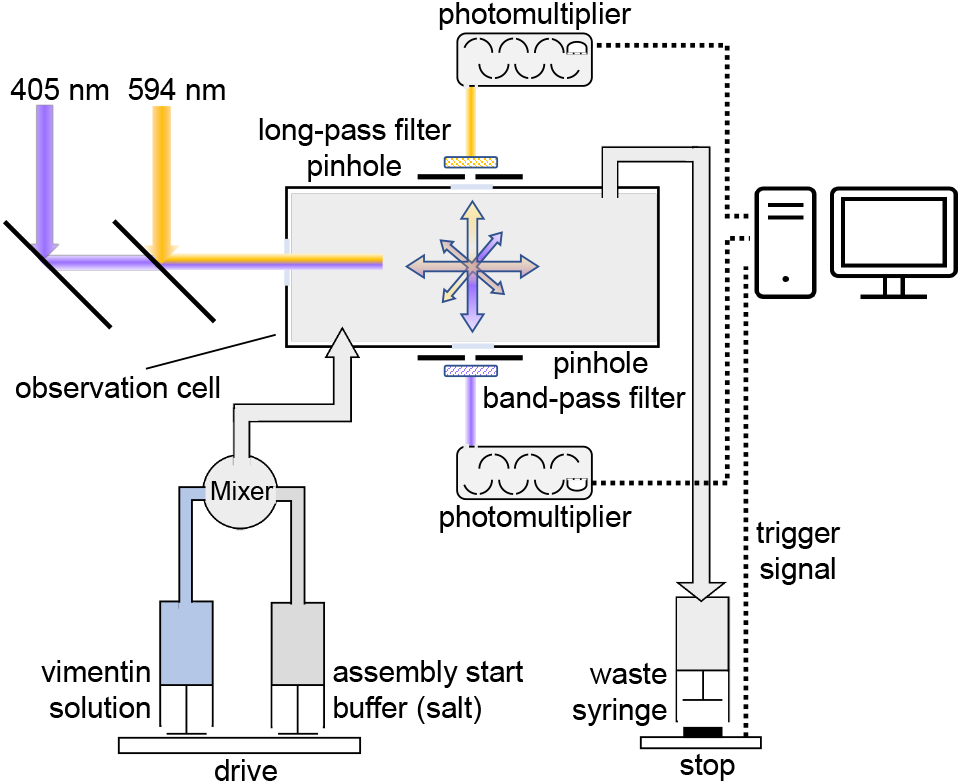
Schematic of the dual laser stopped-flow setup. The aligned 405 nm and 594 nm laser beams are guided into the center of the observation chamber. Perpendicular to the incident beam, the scattered light for both wavelengths is led through a pinhole and a band-pass (405 nm) and long-pass (594 nm) filter, and detected with a photomultiplier.

For measurements, both syringes are emptied simultaneously by a drive platform powered by air pressure at 400 kPa. Equal volumes from both syringes are rapidly (~3 ms) mixed and driven into the detection chamber, displacing the solution from the previous run, which is discarded into the waste syringe. The flow rate of the solution in the system under these conditions is ~1.5 ml/s. The flow is stopped when the plunger of the waste syringe hits the stopping block after 200 *μl*, triggering the data acquisition system. The dead time (delay between trigger activation and signal increase) is 3 ms (8).

Prior to assembly experiments, both syringes are rinsed with several milliliters of de-gassed ddH2O until no change in the light scatter signal is detected. This signal then serves as a baseline for further measurements. In a second step, both syringes are rinsed with several milliliters of tetramer buffer until no change in the light scatter signal is detected. This signal is used to calibrate the light scatter signal. Static light scatter is measured at wavelengths of 594 nm and 405 nm delivered by diode-pumped solid-state lasers (405 nm: Flexpoint 25 mW; Lasercomponents, Olching, Germany; 594 nm: Mambo 25 mW; Cobolt, Solna, Sweden). The light is transmitted via a flexible light guide to the observation chamber. The scattered light is band-pass filtered (405 nm) and long-pass filtered (594 nm) (Schott AG, Mainz, Germany) and detected by two photomultipliers (PM-60s, Hi-Tech Scientific, Salisbury, UK). The electric photomultiplier signals are low-pass filtered (time-constant: 1 ms). Differential signals are digitized (PCI-6023E, National Instruments) at a resolution of 12 bit and sampled at 1 kHz. Measurements that exhibit large fluctuations caused by air bubbles are discarded.

### F. Shape factor calculation

For a solution of randomly oriented rod-like particles such as intermediate filaments with a length > λ/10, when λ is the wavelength of the incident light, and a width < λ/10, Rayleigh-Gans scattering theory considers that the scattered light depends not only on the molecular weight of the scattering object (as in the case of simple Rayleigh scattering), but also on its shape (14). Accordingly, the scattered light intensity *I* at an angle *θ* between the incoming and scattered light direction can be expressed as the scattered light intensity *I*_0_ that would be expected for simple Rayleigh scattering, times a shape factor *P*(*θ*):

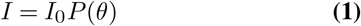

In our setup, the angle *θ* = 90°. In this case, the shape factor for randomly oriented rod-shaped cylindrical particles with length *l* and radius *r* can be computed as the product of a radius-dependent function *F*(*r*) and a length-dependent function *E*(*l*),

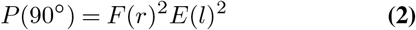

For radii < 10 nm, as is the case for intermediate filaments, the radius-dependent function *F*(*r*) tends to unity. The length-dependent function *E*(*l*) depends on the wavelength λ and can be expressed as

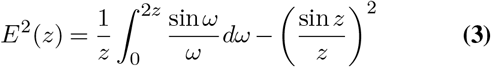

with

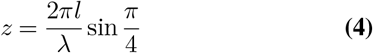

We assume that a single vimentin ULF has a length of 60 nm and that the length increase due to partial filament overlap is 43 nm for each additional ULF (15, 16).

Because the shape factor is wavelength-dependent (Fig. 2A), we can estimate the shape factor and hence the average length of the scattering particles from the ratio of the scattered light intensity measured at different wavelengths (Fig. 2B), if two conditions are met: the shape of the filament length distribution must be known at least approximately (it is log-normal in the case of vimentin), and the average filament length must remain below ~6-7 ULFs, beyond which the scattered light intensity ratio shows a local maximum (Fig. 2B) and little further changes. Importantly, however, the scattered light intensity ratio is largely insensitive to any lateral filament assembly and heterogeneity in thickness, as radial size differences equally affect the scattering signal at different wavelengths.

**Fig. 2.**
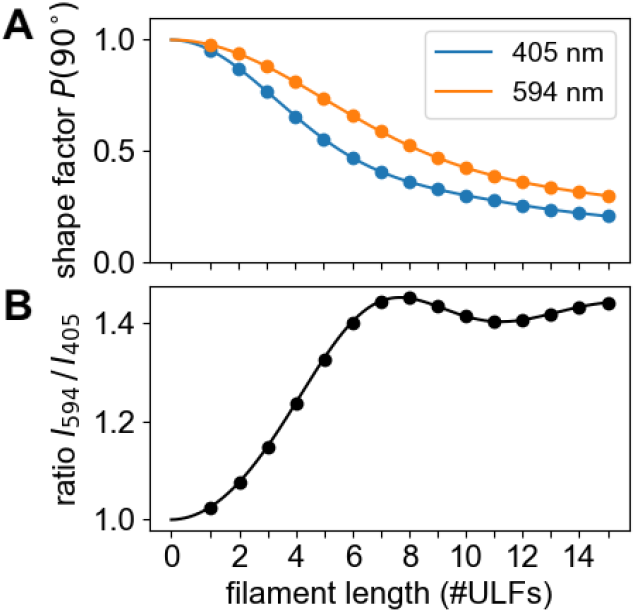
**A)** Shape factor for different wavelengths for a scattering angle of 90°versus filament length as predicted by Rayleigh-Gans theory (Eq. 2-4). The shape factor determines the normalized scattered light intensities for different filament lengths. **B)** The ratio of the scattered light intensities (594 nm / 405 nm) increases monotonically during filament elongation up to a filament length of 7 ULFs.

## Results

### G. Longitudinal filament assembly measured by AFM

Vimentin at a concentration of 0.4 mg/ml is assembled in low salt buffer (50mM NaCl in 25mM Tris-HCl (pH 7.5), and the assembly reaction is stopped by a 10-fold dilution with low salt buffer, followed by fixation by addition of an equal volume of 0.2% glutaraldehyde dissolved in low salt buffer. The filaments are then adsorbed on mica, washed, and dried under a steady stream of nitrogen. AFM measurements (Fig. 3A) confirm a linear increase of filament length with time (Fig. 3B). We find a longitudinal assembly rate of 1.02 ULFs/min. At each time point, the lengths of the individual filaments show an approximately log-normal distribution (Fig. 3C). The width of the distribution increased monotonically with increasing assembly time and hence with increasing average filament length *l***_fil_**. In particular, we find that the geometric standard deviation of the filament length distribution, *σ*, increases monotonically with increasing average filament length (Fig. 3D). This relationship can be empirically expressed with a linear relationship

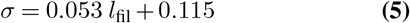

where *l*_fil_ is the geometric mean filament length expressed in units of #ULFs. We have verified using published data that a linear increase of the geometric standard deviation with increasing average filament length can also be observed for assembly of vimentin in KCl salt. These previous measurements indicate that the slope of the relationship may depend on the salt and vimentin concentration, but the intercept remains constant, with values around 0.12.

**Fig. 3.**
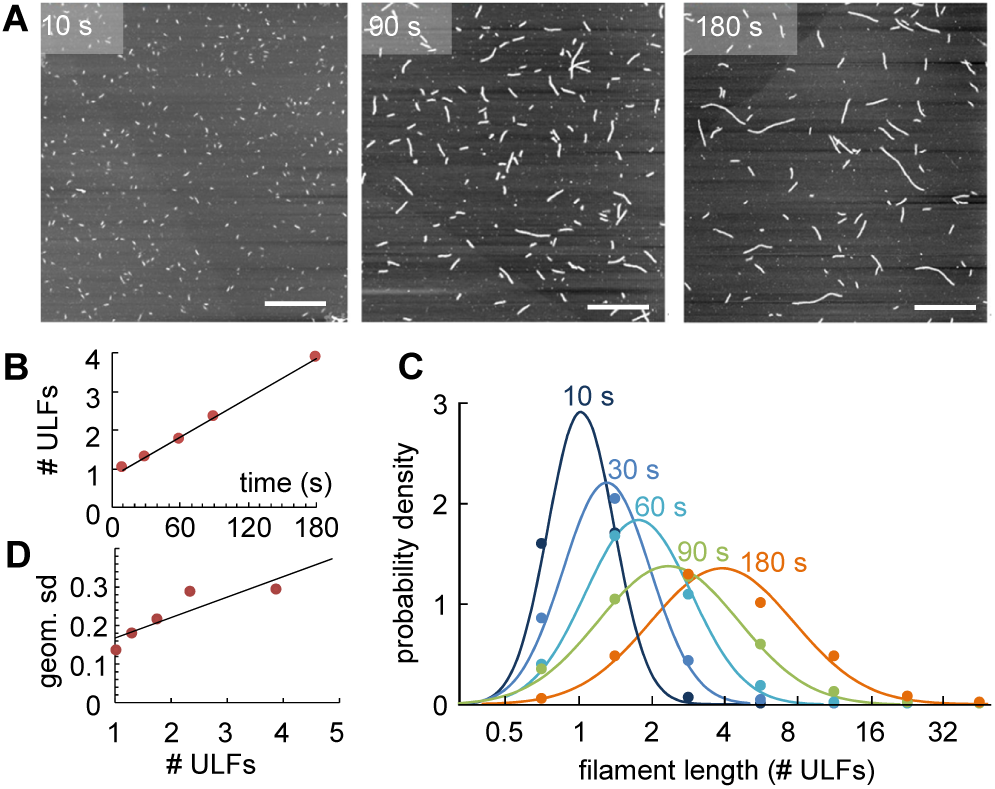
Vimentin elongation measured with AFM. **A)** Vimentin filaments imaged with AFM after 10 s, 90 s, and 180 s after assembly start. Scale bar is 1 *μm*. **B)** Geometric mean of filament length versus assembly time. The black line indicates a linear fit to the data, with a slope of 1.02 ULFs/min. **C)** Distribution of filament lengths at different time points after starting assembly. Points indicate measurements of probability density from AFM images in logarithmically spaced bins (0.5-1 ULF, 1-2 ULFs, 2-4 ULFs etc.); lines are a log-normal fit to the data. **D)** Geometric standard deviation of filament length distribution versus geometric mean filament length. The black line indicates a linear fit to the data, with *σ* = 0.053*l*_fil_ +0.115, when the filament length *l*_fil_ is given in units of #ULFs.

### H. Lateral assembly measured by EM

We use transmission electron microscopy images of vimentin filaments (0.2 mg/ml in 50 mM NaCl) fixated at 2, 10, 30, 60, 180, and 300 s after starting assembly, to measure the filament diameter, which reflects the number and density of tetramers that form the ULFs (Fig. 4A-C). We find that the average filament diameter increases during the first 30 seconds of assembly from 15.5 nm to 17.5 nm, and then decreases to 13 nm during the following 150 s, after which time point the average diameter remains approximately constant (Fig. 4D). This diameter agrees with a previous study (13 nm after 5 min, 12 nm after 10 min, 11 nm after 60 min), where the salt concentration for assembly was 100 mM NaCl (17).

**Fig. 4.**
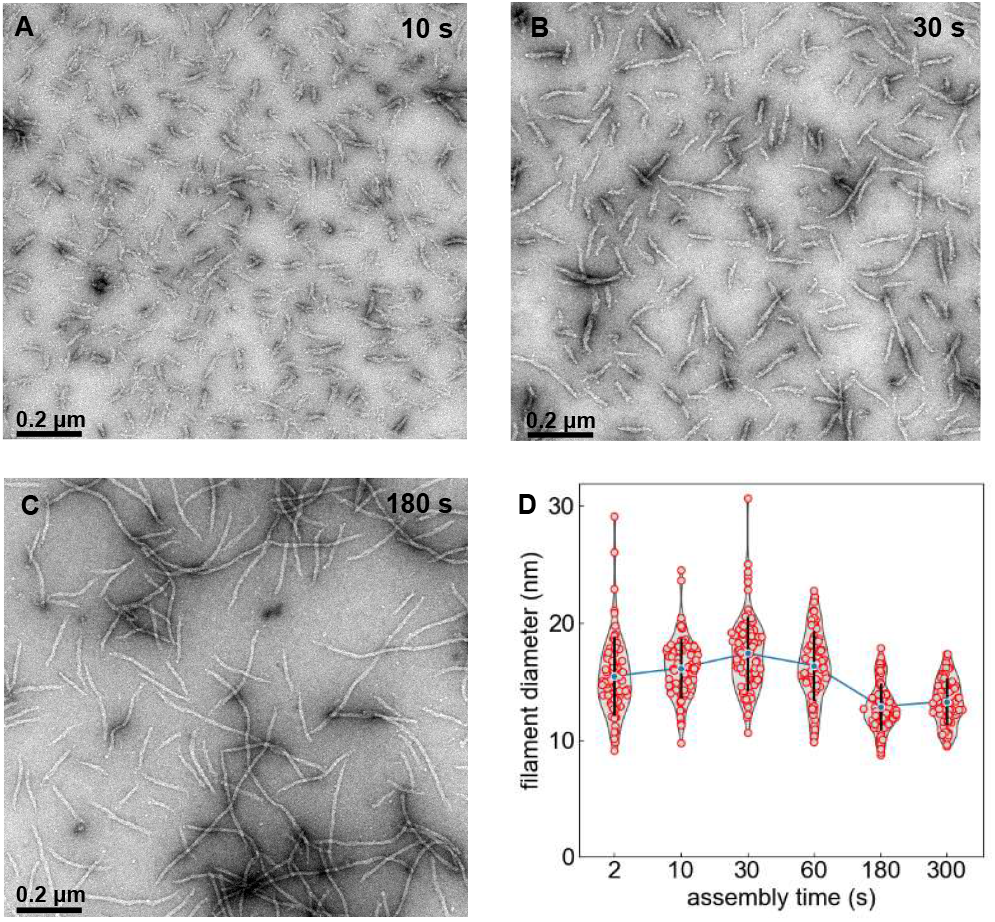
Assembly of vimentin filaments (0.2 mg/ml in 50 mM NaCl) at 37°C for 10 s **(A)**, 30 s **(B)** and 180 s **(C)**. Transmission electron microscopy of glutaraldehyde-fixed and uranyl acetate negatively stained specimens. Scale bar: 200 nm. **D)** Filament diameter and distribution at various time points. Red points represent individual measurements, blue points show the average, black bars denote standard deviation.

Interestingly, the distribution of filament diameters decreases over this time period. While we find some very wide filaments with up to 30 nm diameter during the first 30 s after the start of assembly, not a single filament (from 150 measurements) is found to be wider than 18 nm after 180 s. The mean filament diameter at 30 s of assembly is 30% larger than at 180 s and 300 s. We interpret the reduction in both the mean and the distribution of the filament diameter as the result of an ongoing compaction process, including intra-filamentous reorganization and dissociation reactions of fragments from large ULFs (9).

### I. Longitudinal assembly measured with a dual-wavelength stopped-flow device

We can separate the scatter signal resulting from lateral versus longitudinal assembly by applying Rayleigh-Gans theory. This theory considers the wavelength-dependent decrease of the scattered signal for elongated scatterers relative to the prediction from Rayleigh theory, which assumes point-like scatterers (Fig. 2A). According to Rayleigh-Gans theory, the shape factor for shorter wavelengths decreases more rapidly with increasing filament length, compared to longer wavelengths.

We take advantage of this wavelength-dependency by computing the ratio *r*_*I*594/*I*405_ of the scattered light intensities at 594 nm relative to 405 nm. This ratio only depends on the shape factor and hence on the filament length. Once the filaments have reached a length larger than the wavelength of the incident light, the scatter signal no longer increases as more ULFs are added, and the ratio *r*_*I*594/*I*405_ asymptotically approaches a value of 1.46.

If filament length grows linearly with time, as seen in our data (Fig. 3B), and if all filaments have a uniform length, the ratio *I*_594_/*I*_405_ nm of the scattered light intensities, when plotted versus time *t*, would exactly follow the ratio of the shape factors as shown in Fig. 2B. The effect of log-normal distributed filament lengths, however, blurs the time course of the ratio *r*_*I*594/*I*405_ so that it resembles an exponential function

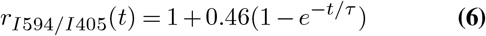

with time constant *τ* as a free parameter. The prefactor 0.46 corresponds to the asymptotic value of the shape factor ratios for infinitely long filaments *P*(90°)_594_/*P*(90°)_405_, minus unity. In practice, we allow this prefactor also to be a free fit parameter, as the experimentally obtained prefactors can range between values of 0.3-0.5. Fit values for the prefactor that are larger than 0.5 indicate in most cases that the measurement was too short to reliably fit a time constant.

Although Eq. 6 provides only an empirical description, it closely describes both a computer-simulated and the experimentally measured time evolution of the ratio *r*_*I*594/*I*405_ (Fig. 5A, 5C). The time constant *τ* of the fit scales inversely with the longitudinal assembly rate *r_la_* (see Fig. 5A inset) according to

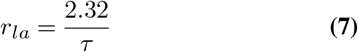

where *r_la_* is given in units of #ULFs/min if *τ* is given in units of min. The factor of 2.32 is determined under the assumption that the relationship between filament length and length distribution that we experimentally determined for 0.4 mg/ml vimentin in low salt (50mM NaCl) conditions (Fig. 3D) also holds for other conditions, which we have not independently verified. Nonetheless, regardless of the specific value for the length distribution, Eq. 6 is still applicable, and the time constant *τ* of the fit scales inversely with the longitudinal assembly rate.

**Fig. 5.**
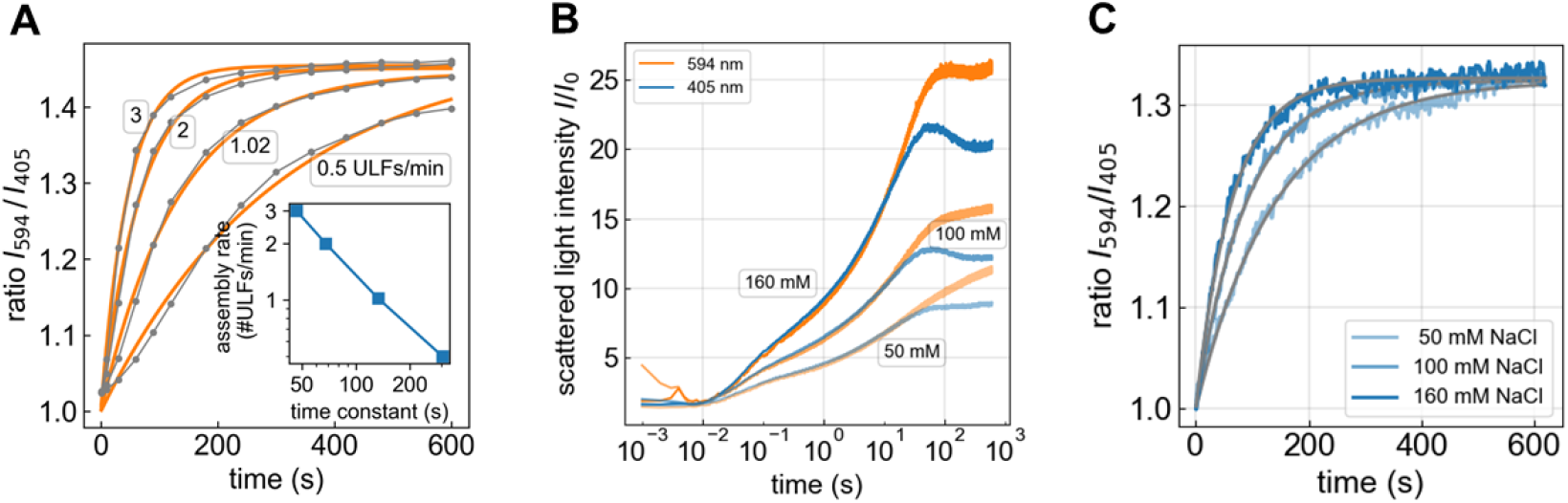
**A)** Monte Carlo simulation of the ratio of the scattered light intensities at 594 nm and 405 nm for different longitudinal assembly rates and time points after starting the assembly (gray curves). The filament length and its distribution versus time is assumed to follow the relationship shown in Fig. 2B and 3D. The orange lines represent the fit of an exponential function (Eq. 6) to the data. The time constant *τ* of the exponential fit scales inversely with the longitudinal assembly rate *r_la_*, according to *r_la_* = 2.32 ULFs/*τ*, if *τ* is given in units of min (inset). **B)** Scattered light intensities at 594nm (orange) and 405nm (blue) over time for assembly of 0.2 mg/ml vimentin in low, medium, and high salt buffer. The scattered light intensities at 405 nm and 594nm develop synchronously but begin to diverge beyond 10 seconds due to the wavelength-dependent scattering properties of elongated filaments. Representative measurements for all other conditions are shown in Fig. SI 2. **C)** Scattered light intensity ratios (594 nm/405 nm) for 0.2 mg/ml vimentin (blue) measured in low, medium, and high salt buffer, with exponential fits (Eq. 6, gray). Representative curves for all other conditions are shown in Fig. SI 2.

We find that the longitudinal assembly of vimentin depends on both, the vimentin and the salt concentration (Fig. 6). For a 4-fold increase in vimentin concentration (from 0.1 to 0.4 mg/ml), we see approximately a doubling of the filament growth rate under medium and high salt conditions, specifically from 1 ULF/min to 2 ULFs/min for medium salt, and from 2 ULFs/min to 4 ULFs/min for high salt. By contrast, under low salt conditions, the filament growth rate increases only slightly with increasing vimentin concentration.

**Fig. 6.**
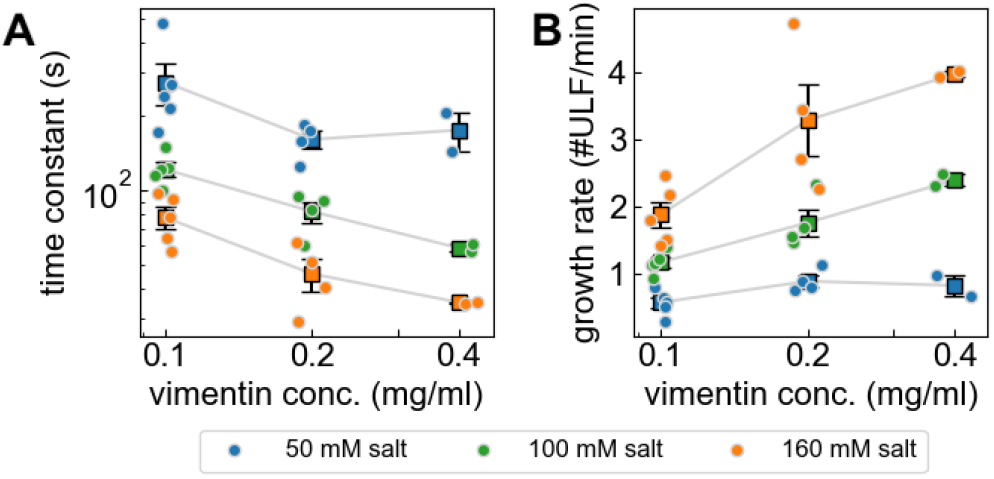
**A)** Time constants of exponential fits (Eq. 6) to the measured 594 nm / 405nm ratios (mean ± SE in black, individual measurements in color) for different vimentin and salt concentrations. **B)** Estimated rate of filament elongation (assembly rate *r_la_*) based on Eq. 7.

### J. Lateral assembly measured with a dual-wavelength stopped-flow device

The time course of the scattered light intensities (Fig. 5B) can be roughly subdivided into three distinct phases: 1) from 10 ms to 1 s after starting the reaction, the scattered light signal is dominated by the lateral assembly of tetramers into ULFs. Longer filaments have not yet formed so that the signals at 405 nm and 594 nm appear identical. The signal intensity rises quickly at first but then, after a first inflection point at around 100 ms, more slowly, without reaching a plateau. Both the speed and the magnitude of the signal intensities increase with increasing salt concentrations. 2) At around 1-5 s, the scattered light signal shows a second inflection and picks up speed, indicating that longitudinal assembly has started to contribute to the scattered light signal. At around 5-30 s, depending on salt concentration, a fraction of the filaments has grown to a length of 2 ULFs or longer, and thus the 405 nm signal falls below the 594 nm signal.

3) The 405 nm signal shows a third inflection point at around 30-60 s (and in the case of medium and high salt concentrations the signals even decrease) before leveling out towards the end of the recording at 600 s. Since the filaments still continue to grow in length, this signal decrease indicates that the filaments undergo a radial compaction. The 594 nm signal, which is less affected by the highly elongated shape of the filaments, shows its third inflection point later at around 100 s, and then decreases only slightly and only at the highest salt conditions before leveling out. The signal plateau indicates that the filaments have reached a length beyond which the scattering signal becomes insensitive to any further growth. The signal plateau, moreover, is considerably higher for higher salt concentrations, indicating that the filaments have a larger mass per cross-section.

Existing models to compute the kinetic rate constants for the initial reaction of tetramers into ULFs (8) do not consider that the diameter of the ULFs may increase at higher salt concentrations. Thus, in the following we do not compute absolute kinetic rate constants but instead describe the influence of salt concentration on the reaction speed relative to the reaction at low salt concentration. We do this by computing the ratio of the scattered light signals, separately for each wavelength, between medium versus low salt conditions (green curves in Fig. 7A,B), and between high versus low salt conditions (orange curves in Fig. 7A,B). These ratios contain information about the lateral and longitudinal assembly kinetics as well as the mass per cross-section of the filaments, relative to the reaction at low salt conditions.

**Fig. 7.**
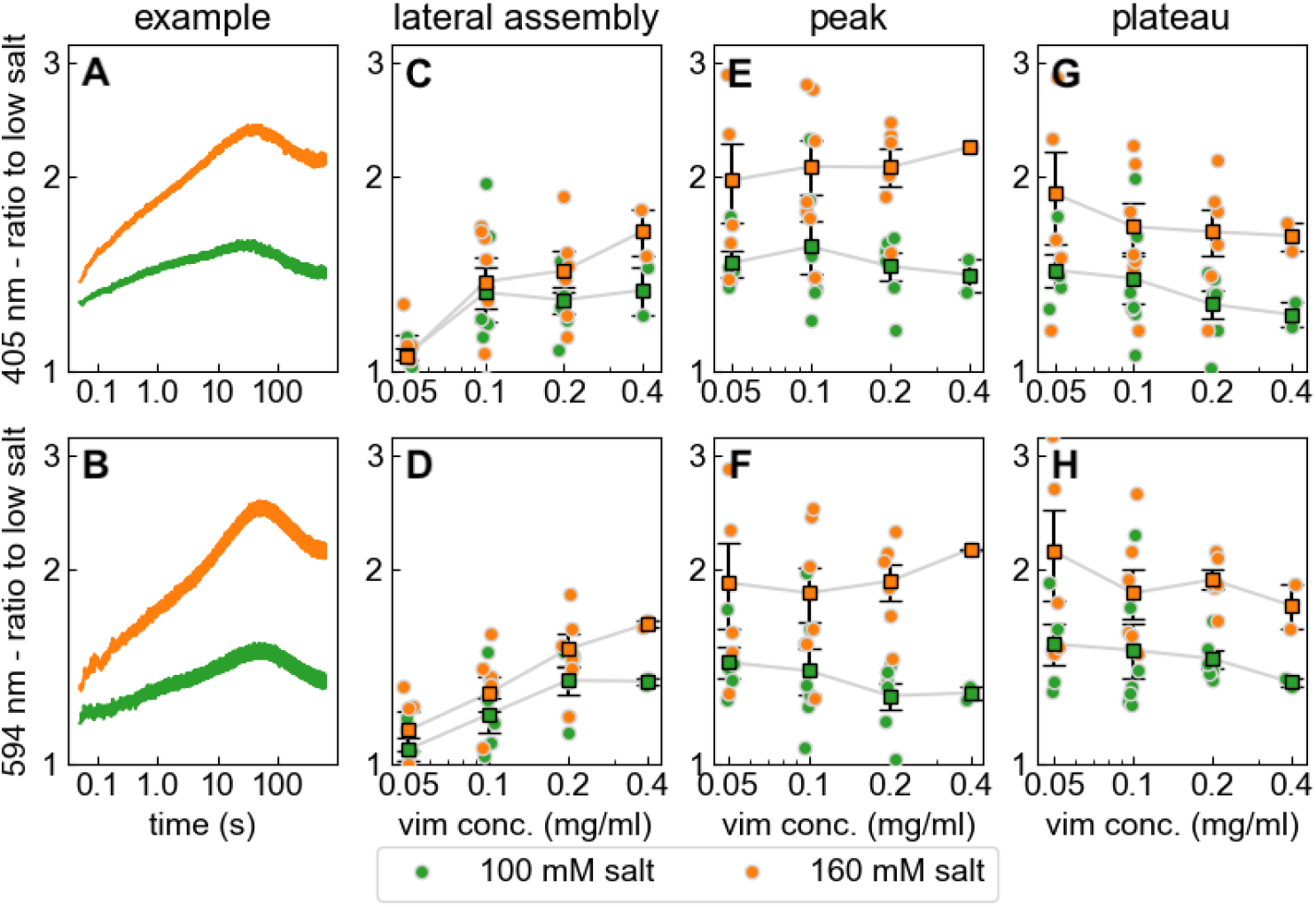
**A,B)** Light scattering signal at 405nm (top row) and 594nm (bottom row) during assembly of 0.2 mg/ml vimentin in 100 mM (green) and 160 mM (orange) salt, normalized by the signal measured for 50 mM salt. **C,D)** Signal ratio (green: 100mM /50 mM; orange: 160mM / 50 mM) as shown in A,B averaged between 20 ms and 40 ms after assembly start, reporting differences in the lateral assembly kinetics relative to the assembly at low salt conditions. **E,F)** Ratio averaged over a 40 s wide interval around the intensity peak, combining contributions from lateral and longitudinal assembly. **G,H)** Ratio averaged between 500-600 s after assembly start, reporting differences in the mass per cross-section of the filaments relative to filaments assembled in 50mM salt buffer.

To decipher this information, we take advantage of the fact that filaments longer than 1 ULF are highly unlikely to have formed in the first 40 ms of the reaction. Thus, the medium-to-low and high-to-low salt ratios, when averaged between 20-40 ms after starting polymerization, report differences in the lateral assembly kinetics. We obtain the same average values for the ratios measured at 405 nm and 594 nm, confirming that longitudinal filament assembly has not yet started. For vimentin concentrations of 0.1 mg/ml and higher, we find on average an approximately 1.3-fold faster assembly under medium salt compared to low salt conditions, and a 1.48-fold faster assembly under high salt compared to low salt conditions. For a vimentin concentration of 0.05 mg/ml, however, the lateral assembly is only marginally faster at medium and high salt compared to low salt conditions. Moreover, we find a trend towards faster lateral assembly (relative for low salt conditions) with increasing vimentin concentration.

We take advantage of the fact that towards the end of the measurement, the filaments have reached a length beyond which the 405 nm scattering signal becomes insensitive to any further growth. The signal ratios between medium-to-low and high-to-low salt conditions, averaged between 500-600 s, therefore report the filament mass per cross-section relative to low salt conditions. We find a 1.3-fold higher mass per cross-section for medium salt, and a 1.6-fold higher mass per cross-section for high salt, compared to filaments that have assembled under low salt conditions. These relative differences are slightly more pronounced for lower vimentin concentrations.

Interpreting the signal ratios for intermediate time points (between 1-100 s) is not as straightforward, as the scattering signal is influenced by both, lateral and longitudinal assembly kinetics. However, since we measure the filament elongation rate for all conditions (Fig. 6), we can, using Rayleigh-Gans theory, estimate the contribution of the elongation kinetics to the scattered light intensities in the absence of any additional lateral growth (Fig. SI 3). Accordingly, the medium-to-low salt ratio of the theoretically predicted scattered light intensities for 405 nm shows a peak at around 30 s after starting the polymerization, with a peak value of 1.15. For the high-to-low ratio, the peak value is 1.2 (Fig. SI3C). The measured peak ratios, by contrast, are considerably higher and reach values of 1.5 and 2.1 for the medium-to-low and high-to-low ratios, respectively (Fig. 7E). The difference between the theoretically predicted and measured values arises from the larger mass per cross-section of the filaments at medium and high salt conditions, compared to low salt. This difference translates to filaments that have a 1.35-fold larger mass per cross-section (for medium salt) and 1.85-fold larger mass per cross-section (for high salt), compared to low salt conditions. These mass per cross-section estimates from the peak ratios are larger than those that we obtain from the plateau ratios (Fig. 7G, 7H). A likely explanation is that the filaments have undergone a maturation process in the form of shedding of loosely associated tetramers, leading to radial compaction. This interpretation agrees with a scanning electron microscopy analysis of filament thickness where we also find that the filament diameter peaks after 30 s of assembly and then decreases by approximately 30 % (Fig. 4D).

## Discussion

Unlike F-actin and microtubules that are formed from globular subunits, intermediate filaments represent a supermolecular self-assembly system that forms linear polymers from fibrous, highly charged molecules (18). The basic architectural units in IFs are parallel coiled-coil dimers with a conserved central *α*-helical domain of about 45 nm in length. The basic functional unit for filament assembly is a tetrameric complex assembled from antiparallel, half-staggered coiled-coil dimers, with the amino-terminal halves of the *α*-helical domain forming a tight interaction. Consequently, this interaction restricts the flexibility of the extended rods significantly (19–22) such that tetramers appear mostly as straight rods in electron microscopic images (4).

When tetramers are induced to assemble laterally into fullwidth unit-length-filaments (ULFs), their interaction is topologically restricted. Thus, tetramers laterally bind each other in register, as concluded from the observation that the length of a tetramer is identical to that of an ULF, i.e. ~60 nm (23). Notably, IFs can be dissociated to tetramers in buffers of low ionic strength, and tetramers will reassemble into filaments when the ionic strength is raised. This property of the IF system allows us to kinetically analyze the formation of filaments from tetramers simply by a change in salt concentration.

The first quantitative kinetic experiments in IFs were performed using electron microscopy (EM) and atomic force microscopy (AFM), by measuring the time-dependent elongation of vimentin filaments (24). Data recorded from 5 s to 20 min after initiating assembly revealed a two-phase scenario: over the first 10 seconds, only ULFs were present. After 30 seconds, first elongation products were observed, and by 60 seconds, the mean filament length was 130 nm, corresponding to the longitudinal annealing of three ULFs. Over the following 20 min, mean filament length increased linearly over time. However, finer details of assembly during the first 60 seconds could not be resolved with this technique. In a previous study, we employed stopped-flow measurements with a time resolution of 3 ms and recorded the assembly of tetramers into the next higher-order complexes such as octamers, 16-mers and ULFs during the first 500 ms (8). However, elongation starts as soon as the first ULFs appear, and hence lateral assembly and elongation occur in parallel and thus cannot be separately measured.

Here, we describe a novel method to simultaneously but separately measure the lateral and longitudinal assembly of intermediate filaments using a dual-wavelength stopped-flow device with a temporal resolution of 3 ms. By observing the static light scattering signal at two distinct wavelengths of 405 nm and 594 nm, we can distinguish between lateral assembly, which increases the signals from both wavelengths by the same factor, from longitudinal assembly, which increases the predominantly the signal from the longer wavelength. The ratio of the 594 nm to 405 nm signal, after subtracting the water background and normalizing the signals during the first 50 ms to the same value, corresponds to the length of rod-like scattering objects if their width is considerably smaller than the wavelength of the incident light, according to Rayleigh-Gans scattering theory. However, if the lengths of the scattering objects are distributed, the average length can only be computed from the 594/405 ratio if the shape of the length distribution is known. From AFM measurements, we know that the filament lengths are log-normal distributed, and that the width of the distribution, apart from a constant offset, increases linearly with the geometric mean of the filament length (Eq. 5). In practice, we take advantage of the observation that vimentin filaments elongate with a constant assembly rate, and that the 594/405 ratio increases according to a mono-exponential function (Eq. 6). Hence, the time constant of the exponential function is inverse proportional to the longitudinal assembly rate, according to Eq. 7.

The factor of proportionality in Eq. 7 is subject to several uncertainties. First, we assume that the assembly rates and filament length distribution in the AFM and stopped-flow experiments are similar. AFM measurements require fixation of growing filaments and their attachment to a solid support, which has been argued to be a potential source of error in determining the length and diameter of filaments (25, 26). However, we demonstrated, by performing EM and AFM on different substrates, that both methods produce highly consistent results (10, 24). Second, we further assume that the filament length distribution depends only on the geometric mean of the filament lengths but not on salt or protein concentration. We have not tested this assumption experimentally, but a recent modeling study confirms its validity (27).

We apply our method to investigate the assembly kinetics of vimentin and its dependence on the ion concentration of the assembly buffer. Our data are in line with earlier stopped-flow and electron microscopy studies: the early assembly during the first second is dominated by lateral reaction processes. Between 1-10 s, longitudinal assembly starts to dominate, which can be observed by the differences in the scattered light intensities at different wavelengths. Lateral and longitudinal assembly processes continue to coexist for the next 10-30 s. This is followed by a phase from about 30-90 s where the filaments continue to grow in length and - dependent on salt and vimentin concentration - shrink in diameter. This reduction in diameter is seen both by electron microscopy measurements (Fig. 4) and by the overshoot in the scattered light intensities at 405 nm (Fig. 5B). The reduction in diameter is facilitated by a dissociation of smaller complexes from the filament core and by intra-filamentous reorganization, which is also referred to as radial compaction (9, 17). After several minutes, the lateral compaction process comes to a halt, and longitudinal assembly prevails.

We find a strong salt concentration-dependent increase of both, the lateral and longitudinal assembly kinetics. Notably, we find that filaments assembled at higher salt concentrations (100 and 160 mM) exhibit a greater mass per cross-section and show excess radial growth during the first 30-90 s of assembly, compared to assembly at 50 mM salt. At higher salt concentrations, the kinetics of lateral association is speeded up to an extent that heterogeneous ULFs are formed, containing regular 8 to 12 tetramers that are productive for elongation, together with hyper-aggregated assembly complexes that are unstable. Such heterogeneous hyper-aggregates have previously been observed by scanning transmission electron microscopy (STEM) when filament assembly was triggered by salt addition (9). By contrast, when filaments were assembled by dialysis, IFs were completely homogeneous, with a constant mass distribution along individual filaments (4). This, together with our observation that the signal overshoot during the first 30-90 s of assembly at a lower salt concentration of 50 mM is largely absent, indicates that an ordered lateral assembly needs a certain time for the precise association of tetramers. Notably, different IF proteins vary considerably in their assembly kinetics: keratin K8/K18 assembles about a hundred times faster than vimentin, and the sequence-related desmin assembles five times faster than vimentin (28). Hence, despite the principally identical structural organization and assembly mechanism of these different IF proteins, their individual amino acid sequences appear to have a strong impact on the kinetics of the longitudinal assembly reaction. However, the minimum time required for an orderly, precise lateral association of tetramers into ULFs is not reduced to the same extent.

Our findings are in line with a recent report by Lopez and colleagues who have demonstrated, using static and dynamic light scattering in combination with mass measurements by scanning transmission electron microscopy, that the ionic strength of the assembly buffer influences both the lateral and the longitudinal kinetics of assembly. To avoid problems arising when IF proteins are assembled under near-physiological conditions, such as very fast formation of filaments in combination with the formation of extensive filament bundles (29), they employed a low protein concentration (0.035 mg/ml) and an assembly temperature of 20°C (25, 26). In such a regimen, the highly active molecular interactions of IF tetramers and the formation of higher order assembly products are drastically decelerated as compared to assembly at 37°C and higher protein concentrations (8). Within 120 minutes of assembly, they obtained a mean filament length representing four longitudinally annealed ULFs. For comparison, under the conditions employed in our experiments (protein concentration 0.05-0.4 mg/ml, assembly at 37°C), a median length of four ULFs is reached within 1-4 min (Fig. 6). Lopez et al. also noted that at higher salt concentrations, the mass per cross-section of the filaments increased (26). Furthermore, the filament growth rate decreased with increasing filament length *L* according to *L*^−4^. By contrast, our results obtained at 37°C and at higher protein concentrations demonstrate an approximately constant, length-independent growth rate, giving rise to a nearly linear filament growth over time (15).

We have previously demonstrated that vimentins from different species exhibit a temperature-dependent assembly such that above a certain temperature (usually the body temperature of these organisms), the longitudinal assembly is impaired, and instead band-like fibrous aggregates appear (4, 30). Also at low temperature, the formation of loosely associated filaments is prominent but elongation is impaired, as demonstrated for amphibian vimentin, which assembles best at 28°C but fails to regularly assemble at 4°C (31). Hence, it is not clear in which temperature range vimentin, of a certain species, is assembling in a regular manner. This question can of course now be tackled with our stopped-flow system.

In vivo, the longitudinal assembly is spatially and kinetically governed by a chaperone system involving motor protein-mediated distribution of ULFs, thereby controlling firstly the subunit composition of ULFs, and secondly the productive elongation within the cell (32). Notably, a bifurcation of filaments is not observed, neither in vitro or in vivo. An ULF segment harboring, for instance, 12 tetramers could theoretically connect to two filaments with 6 tetramers each. This has, however, never been observed by us in hundreds of EM and AFM micrographs, indicating that, even if such an event would take place, the resulting structure would not be stable.

The salt concentration-dependent speed-up of assembly is likely caused by the efficient shielding of ionic charges between the ionic amino acid side chains within and between dimers. Hence, with more monovalent ions being present, equal ionic charges are hindered from repelling each other, and salt bridges may form more readily. As a result, organized interactions leading to productive assembly may proceed more efficiently. This is of great importance for disease mutants of both vimentin and desmin. They have previously been described to leave the regular assembly pathway at distinctly different time points (13, 33–35). Accordingly, we predict that assembly of such mutated IFs requires higher salt concentrations compared to wildtype IFs. Indeed, the desmin mutant R406W-desmin, which causes severe myopathies and cardiomyopathies in humans, was characterized to assemble not much beyond the ULF state when assembly was conducted in 50 mM sodium chloride (13). By contrast, when assembly was conducted at 160 mM sodium chloride, R406W-desmin exhibited the same assembly kinetics as WT-desmin during the first 30 seconds of assembly. By this time, however, the nascent filaments agglomerated laterally, thereby blocking their longitudinal annealing. As a result, assembly terminated with the accumulation of unproductive non-IF aggregates (12).

## Conclusion

The measurement of static light scattering at two different wavelengths in a stopped-flow device allows for the calculation of the shape factor of the reaction product. This in turn can be used to infer the kinetics of both the lateral and longitudinal assembly of intermediate filaments with unprecedented temporal resolution under a wide range of conditions, such as different temperatures and ionic strengths, the presence of chaperones, cross-linkers, or reactive oxygen species.

## Author Contributions

NM conceived the method and modified the stopped-flow device for dual-wavelength measurements, NM, HH, WHG and BF designed the experiments, NM, HH, BF, WHG, and LS carried out the experiments, LS, BF, and NM analyzed the data, SP performed the simulations, NM, HH, BF and LS wrote the manuscript.

## Acknowledgements

We thank Kati Toth (B040, DKFZ) for stimulating discussions regarding Rayleigh-Gans theory, Werner Schneider for help with setting up the dual-wavelength stopped flow device, Lisa Kamm for help with generating the recombinant vimentin, and Dorothea Schultheis for help with immunofluorescence microscopy. This work was supported by the German Research Foundation (DFG, HE1853/15-1) and a Discovery Grant of the Natural Sciences and Engineering Research Council of Canada (RGPIN-2018-04967).

## Supplementary Note 1: Figures

**Fig. SI 1.**
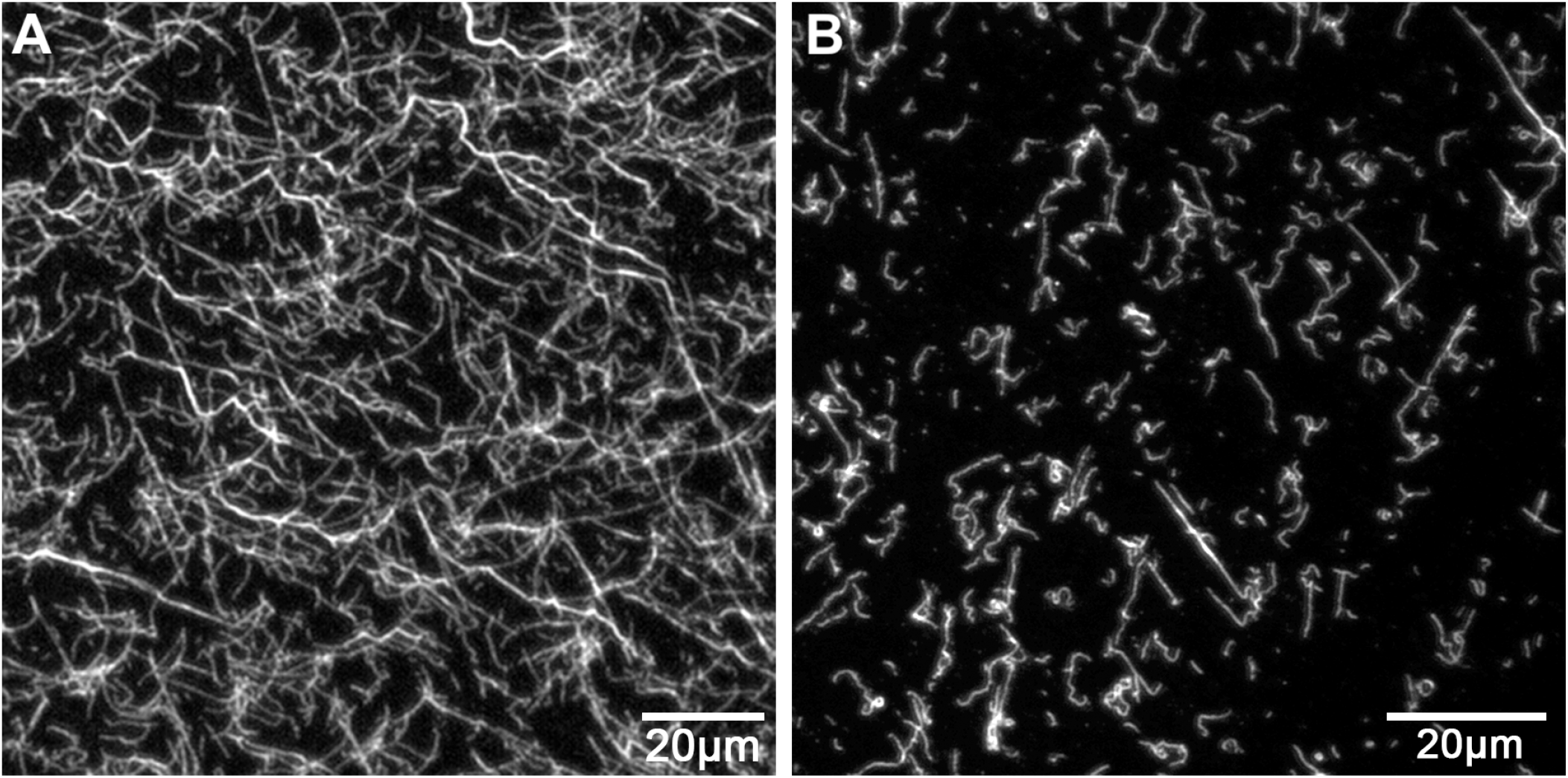
Immunofluorescence microscopic images of vimentin filaments after 60 min of assembly (0.2 mg/ml, 160 mM NaCl). **A)** The filaments form a complex network. **B)** In a 1:50 diluted suspension, single unbranched and highly flexible filaments are visible. Scale bar: 20 *μm*.

**Fig. SI2.**
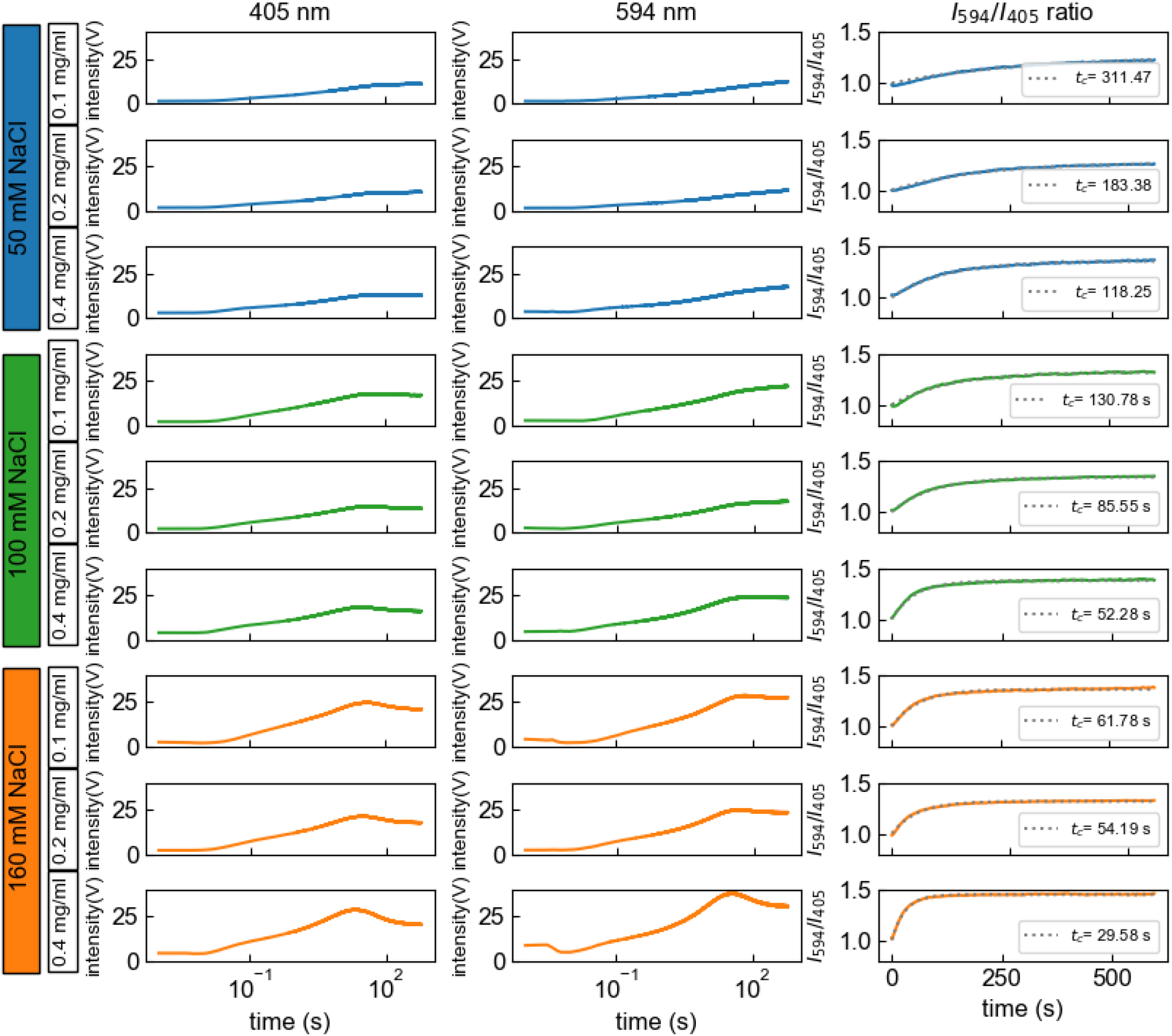
Normalized mean scattered light intensities at 405 nm and 594nm and their intensity ratio for NaCl concentrations of 50 mM (blue), 100 mM (green) and 160 mM (orange), and for vimentin concentrations of 0.1 mg/ml, 0.2 mg/ml and 0.4 mg/ml. The time constant *t_c_* of the exponential fit of the intensity ratio signal, as displayed in each figure, decrease with increasing vimentin and salt concentration.

**Fig. SI3.**
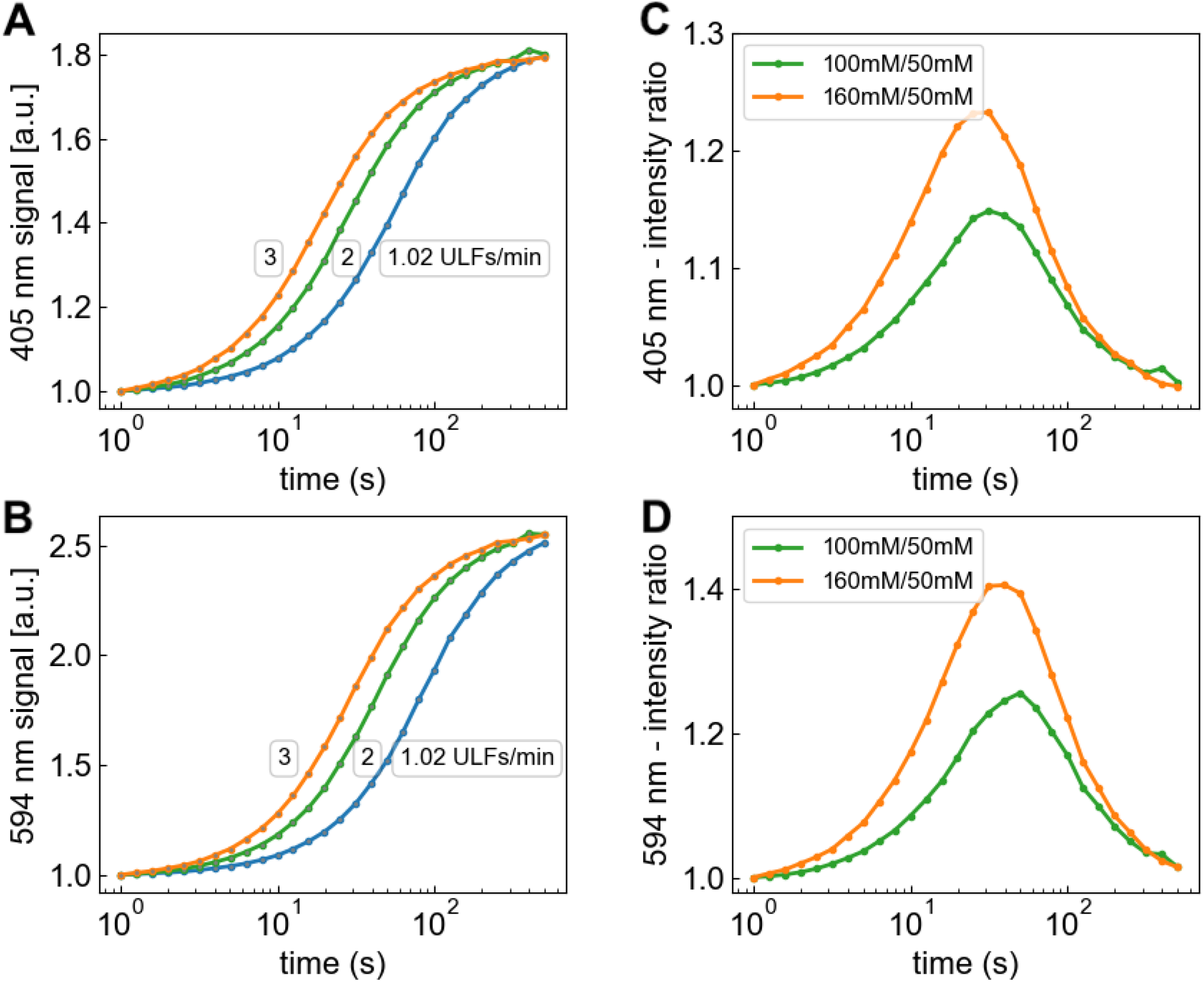
Simulated scattering intensity during longitudinal assembly **(A,B)** for growth rates 1.02 (blue), 2.0 (green) and 3.0 ULFs/min (orange). These growth rates correspond to the assembly of 0.2 mg/ml vimentin in 50 mM (blue), 100 mM (green) and 160 mM (orange) sodium chloride, respectively. The simulated scattered light curves for 100 mM (green) and 160 mM (orange) sodium chloride are divided by the scattered light curve of 50 mM (blue) NaCl **(C,D)**. The different assembly speeds for each salt condition result in a peak in the 100 mM / 50 mM (green) and the 160 mM / 50 mM (orange) signal ratios between 30 s and 90 s, but beyond 500 s, the ratios return to unity as the filaments under all conditions have elongated to a length of >10 ULFs beyond which the scatter signal shows little or no further increase. The measured signal ratios (Fig. 7), by contrast, do not return to unity, indicating that the mass per cross-section of the filaments increases with increasing salt concentrations.

